# Differential roles of Rad18 in repressing carcinogen- and oncogene-driven mutagenesis *in vivo*

**DOI:** 10.1101/2025.06.30.662411

**Authors:** Jay R. Anand, Bethany Wagner Brown, Jitong Lou, Qisheng Gu, Yang Yang, Gaith Droby, Di Wu, Akankshya Jena, Jialiu Xie, Yuri Fedoriw, Bernard Weissman, Scott Williams, Cyrus Vaziri

**Affiliations:** Department of Pathology and Laboratory Medicine, University of North Carolina at Chapel Hill, Chapel Hill, NC 27599, USA; Department of Biostatistics, University of North Carolina at Chapel Hill, Chapel Hill, NC 27599, USA; Division of Oral and Craniofacial Health Sciences, University of North Carolina at Chapel Hill, Chapel Hill, NC 27599, USA; Curriculum in Genetics and Molecular Biology, University of North Carolina, Chapel Hill, NC 27599, USA; Lineberger Comprehensive Cancer Center, University of North Carolina at Chapel Hill, Chapel Hill, NC 27599, USA

## Abstract

The DNA repair protein RAD18 activates ‘Y-family’ Trans-Lesion Synthesis (TLS) DNA polymerases that are DNA damage-tolerant and potentially error-prone. RAD18 is also frequently overexpressed and pathologically activated in cancer cells. However, the extent to which RAD18 shapes cancer genomes and impacts tumorigenesis is unclear. Therefore, we tested the effect of *Rad18* status on chemically-induced and oncogene-driven tumorigenesis. In a chemically-induced oral carcinogenesis model, acute (2-16 days) 4NQO-treatment induces expression of *Rad18* and TLS polymerase mRNAs in mouse oral epithelial cells prior to emergence of oral squamous cell carcinomas (OSCCs). Chronic (8 week) 4NQO-treatment leads to onset of oral tumors that is accelerated in *Rad18^-/-^* mice when compared with *Rad18^+/+^* animals. Analysis of OSCC exomes reveals increased levels of G(C)>T(A) transversions in *Rad18^-/-^* tumors when compared with *Rad18^+/+^*. Therefore, Rad18 promotes error-free bypass of 4NQO-induced DNA lesions and suppresses 4NQO-induced oral carcinogenesis. In a *Kras^G12D^*-induced lung carcinogenesis model, *Rad18*-deficiency did not affect rates or incidence of oncogene-induced lung tumors or mutations. Taken together, we demonstrate that Rad18 has context-specific tumor-suppressive activity. Given the prevalence of 4NQO-like environmental exposures, RAD18 is highly likely to shape human cancer genomes and perhaps influence other aspects of the tumorigenic process.

## INTRODUCTION

DNA damage and DNA repair are intimately connected with all phases of cancer. Carcinogenesis is often initiated by environmental genotoxic exposures (e.g. solar radiation, tobacco smoke). Error-prone replication or repair of damaged DNA can induce mutations in growth-regulatory genes and incite multistep carcinogenesis (1–3). The efficiency and fidelity with which carcinogen-induced DNA damage is repaired directly impacts mutagenic and tumorigenic endpoints. Deployment of error-prone DNA repair mechanisms in lieu of high-fidelity processes can lead to cancer-causing mutations such as Single Nucleotide variants (SNVs) as well as genomic structural variants (SVs, including large deletions, translocations, and complex rearrangements). Therefore, DNA repair pathway availability and DNA repair pathway choice can be highly consequential.

The action of every error-prone DNA repair process on each species of DNA lesion generates a hallmark mutation.(4–7) Cancer cells typically have unique ‘mutational portraits’ that reflect the history of all the error-prone DNA replication and repair events that occurred during multi-step carcinogenesis (4,5). In the last decade, the ‘mutation signature’ technique has emerged as a convenient way to annotate and curate the individual mutational patterns most commonly observed in human cancer (http://cancer.sanger.ac.uk/cosmic/signatures) (8). Using the mutation signature technique, different Single Base Substitution (SBS) Signatures (representing single nucleotide variants or SNVs) are annotated based on sequence context. For any given SNV, there are 96 context-dependent mutation frequencies that consider the two flanking 5’ and 3’ nucleotides (8). The 30 different SBS signatures are defined by the frequencies with which each SNV occurs in all possible trinucleotide settings. Additionally, Doublet Base Substitution (DBS) signatures may be extracted based on concurrent alterations in two consecutive bases. Small insertion and deletion (ID) signatures can be annotated based on gain or loss of small (1-50 bp) DNA fragments. Finally, Copy Number Variations (CN) signatures can be determined based on copy number profiles. There is immense interest in identifying the mechanisms responsible for different mutational signatures in cancer cells. Elucidating molecular underpinnings of mutational signatures will reveal key events that drive cancer, predict responsiveness to therapy and provide opportunities for precision medicine. Remarkably, the underlying mechanisms responsible for approximately one third of mutation signatures are still unknown (9). To definitively reveal the molecular etiology of mutational signatures it is necessary to experimentally determine how specific combinations of genotoxins with DNA repair defects shape the genomic landscape.

In addition to generating the mutations that drive cancer, DNA repair processes help sustain pre-neoplastic and neoplastic cells by conferring DNA damage tolerance. Cells undergoing multi-step carcinogenesis exist in stressful environments and experience DNA damage from intrinsic sources. Activated oncogenes are a major source of intrinsic genotoxicity in pre-neoplastic and neoplastic cells (10). Oncogenes induce DNA replication stress and DNA damage via multiple mechanisms. For example, oncogenes can dysregulate DNA replication licensing leading to re-replication and head-to-tail replication fork collisions (11,12). Additionally, oncogenes can stimulate transcription and increase replication-transcription conflicts (RTCs). Oncogenes also disrupt R-loop homeostasis which further promotes RTCs (13). Oncogene signaling can alter nucleotide metabolism and limit concentrations of dNTP precursors available for DNA synthesis - ultimately causing DNA replication stalling and DNA damage (14). Some oncogenes also stimulate ROS production and cause oxidative DNA damage (15,16). Therefore, genome maintenance processes that allow neoplastic cells to remediate or tolerate the inherent genotoxic stresses associated with tumorigenesis may promote cancer cell viability. Clearly, mutagenesis and DNA damage tolerance are enabling characteristics of cancer cells. Therefore, identifying the genome maintenance pathways that drive mutagenesis and confer DNA damage tolerance in neoplastic cells will explain fundamental mechanisms of carcinogenesis and might reveal critical dependencies and new therapeutic targets.

‘Trans-Lesion Synthesis’ (TLS) is an appealing candidate mechanism of mutagenesis and DNA damage tolerance during multi-step carcinogenesis. TLS uses error-prone ‘Y-family’ DNA polymerases to replicate or repair DNA templates containing diverse species of DNA lesions. Therefore, TLS can confer DNA damage tolerance at the expense of DNA replication/repair fidelity (17,18). Activation of the canonical TLS pathway, and recruitment of Y-family TLS polymerases to sites of DNA damage is regulated by ubiquitin signaling (19). During S-phase, encounters between replicative DNA polymerases and bulky DNA adducts (or other hard-to-replicate DNA templates) lead to DNA replication fork stalling. The E3 ubiquitin ligase RAD18 is recruited to stalled DNA replication forks and mono-ubiquitinates the polymerase processivity factor PCNA at a conserved lysine residue, K164 (19,20). Mono-ubiquitinated PCNA then helps recruit the Y-family DNA polymerases eta (Polη), kappa (Polκ), iota (Polι), and REV1 to the vicinity of stalled replication forks (17,21,22). TLS polymerases have specialized ubiquitin-binding zinc fingers or ubiquitin-binding motifs (termed UBZ and UBM domains respectively) which mediate their stable engagement with the stalled replication forks (23). The TLS polymerases have open catalytic clefts and active sites that can accommodate damaged DNA templates (17). Collectively, the Y-family TLS polymerases allow cells harboring damaged genomes to maintain S-phase progression. However, TLS polymerases are inherently less accurate than replicative DNA polymerases (Polε and Polδ). Therefore, deploying TLS polymerases for lesion bypass carries risk of mutagenesis.

The extent to which TLS is error-free or mutagenic depends on the nature of the DNA lesion and the Y-family DNA polymerase(s) selected for replicative lesion bypass. Some TLS polymerases have preferred ‘cognate’ DNA lesions that they can replicate in a relatively error-free fashion (17). For example, Polη is able to replicate DNA templates harboring its cognate lesions, solar UV-induced cyclobutane pyrimidine dimers (CPD) with accuracy (17). However, when Polη is absent, compensatory error-prone TLS by Polκ and Polι can lead to mutations (24). During S-phase, TLS polymerases can enable a stalled leading strand to continue synthesizing DNA (a process termed ‘TLS on-the-fly’). TLS polymerases can also fill the ssDNA gaps that arise when repriming generates a new replication fork downstream of a stalled DNA polymerase (25). In addition to rescuing stalled replication forks and eliminating post-replicative discontinuities, TLS polymerases can contribute to filling of single-stranded DNA gaps that arise in G1 and G2/M as a byproduct of other genome maintenance processes such as NER, BER and MMR (26–29).

Several lines of evidence may suggest a role for RAD18-mediated TLS in promoting tumorigenesis. For example, some COSMIC mutational signatures are attributed to Y-family DNA polymerases (30–32), indicating that error-prone TLS occurs as normal cells undergo transformation and malignant progression. RAD18 and TLS polymerases are necessary for primary cells to tolerate ectopically-expressed oncogenes (33), suggesting that TLS is required viability of cancer cells. Indeed, many cancer cells are sensitive to TLS inhibition (34).

Consistent with a dependency of many cancer cells on TLS, *RAD18* mRNA is expressed at high levels in most tumors when compared with adjacent normal tissues (22). The RAD18 protein is also pathologically stabilized by the Cancer/Testes Antigen (CTA) MAGEA4 in many cancer cells (35), also suggesting that RAD18 helps neoplastic cells sustain tumorigenic phenotypes.

Although RAD18-mediated DNA repair mechanisms have been extensively studied, and RAD18 overexpression and pathological activation in cancer cells is well-established, remarkably little is known regarding its impact on carcinogenesis. In a study using the carcinogen DMBA to induce skin carcinogenesis, Rad18 promoted DMBA signature A>T mutations (36). Furthermore, *Rad18*-deficiency led to a distinct DMBA-induced mutational signature, but did not affect incidence of tumor formation when compared with wild-type mice. In a separate study, the incidence of UV-induced skin tumors was indistinguishable between *Rad18^+/+^* and *Rad18^-/-^*mice (37). Therefore, Rad18 may have pleiotropic roles in cancer that are highly context-dependent.

Here we use well-characterized experimental models to test the roles of Rad18 in chemically-induced oral carcinogenesis and oncogene-induced lung tumorigenesis. 4-nitroquinoline-1-oxide (4NQO) is a DNA damaging agent that is widely used to induce OSCCs that are pathologically similar to human OSCC. Moreover, mutations generated by 4NQO show remarkable similarity with those observed in human OSCCs (38,39). TLS is implicated in tolerating 4NQO-induced DNA damage(40–42) and the TLS inhibitor JH-RE-06 sensitizes cells to 4NQO (43). However, previous studies have not tested the contribution of TLS to oral carcinogenesis and mutagenesis. Here we show that expression of *Rad18* and Y-family TLS polymerases is dynamically regulated at the mRNA level during preneoplasia. Our analysis of TCGA patient cohorts also shows that both *RAD18* and its pathological activator *MAGEA4* are over-expressed in human HNSCC. Surprisingly, we demonstrate that *Rad18*-deficient mice show increased rates of mutagenesis and oral carcinogenesis when compared with wild-type animals. Conversely, MAGEA4 overexpressing mice also show resistance to 4NQO-induced carcinogenesis. Unexpectedly, Rad18 does not impact *Kras*-driven lung tumorigenesis, even although TLS is required for cells to tolerate Kras-induced DNA replication stress (33). Our genomic analyses show that Rad18 also differentially impacts the mutational landscapes of 4NQO- and Kras-induced tumors. Taken together, we demonstrate an unexpected new tumor-suppressive role for Rad18. We also show that depending on the cancer-causing agent, Rad18 can have context-specific effects on tumorigenesis and the cancer genome.

## MATERIAL AND METHODS

### Cells and culture

Human oral keratinocytes immortalized with hTERT (OKF-6) were gift from Dr. Natalia Isaeva at UNC and the dysplastic oral keratinocyte (DOK) cell line were obtained from Millipore Sigma (DOK, 94122104). OKF-6 cells were cultured in Keratinocyte SFM (serum-free medium) (Gibco, 17005042) supplemented with KGM Singlequot Supplementary Pack (Lonza, CC-4151), streptomycin sulfate (100 μg/ml) and penicillin (100 units/ml). While the DOK cells were cultured in Dulbecco’s Modified Eagle’s Medium (DMEM) supplemented with 2mM Glutamine, 5ug/ml hydrocortisone, and 10% fetal bovine serum, streptomycin sulfate (100 μg/ml) and penicillin (100 units/ml).

### SDS-PAGE and immunoblotting

To induce DNA damage, cells were reverse transfected with siControl and siRAD18 for 48 hours and then were treated with UV (20 J/m2) or 4NQO (0.01 μM). Four hours post-treatment, chromatin or soluble extracts were prepared as previously described.(44) Briefly, cells were washed three-times with ice-cold phosphate buffered saline (PBS) and lysates were prepared with 200ul ice-cold cytoskeleton buffer (CSK buffer; 10 mM PIPES, pH 6.8, 100 mM NaCl, 300 mM sucrose, 3 mM MgCl2, 1 mM EGTA, 1 mM dithiothreitol, 0.1 mM ATP, 10 mM NaF, and 0.1% Triton X-100) freshly supplemented with complete protease inhibitor cocktail (Roche, Indianapolis, IN, USA) and PhosSTOP (Roche). Chromatin-enriched extracts were generated by centrifuging the lysates at 1500 × *g* for 4 min and removing the CSK-soluble fraction. The CSK--insoluble chromatin-enriched fractions were washed with 1 ml of CSK buffer, resuspended in 100ul CSK containing nuclease. Chromatin-enriched extracts were studied by sodium dodecyl sulphate-polyacrylamide gel electrophoresis (SDS-PAGE) and immunoblotting as described previously. The following primary antibodies were used for immunoblotting analysis: p-ATM S1981 (Santa Cruz Biotechnology, sc-47739), mouse monoclonal PCNA clone PC10 (Santa Cruz Biotechnology, sc-56), RAD18 (Bethyl Laboratories Inc., A301-340A), GAPDH (Santa Cruz Biotechnology, sc-32233), phospho-Histone H2A.X (Ser139) (Millipore Sigma, 05-636), β-actin (Santa Cruz Biotechnology, sc-130656), phosphor-CHK1 (Cell Signaling Technology, 12302), and rabbit polyclonal anti-H3 (9715, Cell Signaling Technology). Perkin Elmer Western Lightning Plus ECL was used to develop films.

### Viability assay

DOK cells were seeded at densities of 2000 cells/well in triplicate in 6-well plates. Relevant concentrations of 4NQO were prepared by diluting it in the growth medium from DMSO-based stock solutions. These prepared concentrations in growth medium were then introduced directly to the cells. After a period of 3 days, the cell was replenished with growth medium containing fresh drug. To observe colonies, culture plates were treated with a solution of 0.05% crystal violet in 1× PBS with 1% methanol and 1% formaldehyde. Scanning of the plates was done utilizing an HP scanner. For clonogenic survival assessments, the stained plates were scanned using HP scanner, and the ImageJ plugin ColonyArea was employed to automatically measure the colony quantities. Statistical analysis was performed using two-way analysis of variance (ANOVA) with Sidak’s multiple comparisons test.

### Mice and breeding

Mice were maintained in an AAALAC accredited facility on a diet of standard chow. All experiments were carried out in accordance with the Institutional Animal Care and Use Committee (IACUC)-approved animal protocols. *RAD18^+/-^* mice on a C57BL/6J genetic background were gifts from Dr. Satoshi Tateishi and have been previously described (45,46). *LSL-K-ras^G12D^* mice (gifts from Dr. Tyler Jacks) and *p53^fl/fl^* mice (gifts from Dr. Aton Berns) mice on a C57BL/6J genetic background have been previously described (47,48). *LSL-K-ras^G12D^*, *p53^fl/fl^* and *Rad18^+/-^* strains were crossed to generate the *Rad18^+/+^* control *LSL-K-ras^G12D^*; p53^fl/fl^ mice and *LSL-K-ras^G12D^*; *Rad18^+/-^*; p53^fl/fl^ mice. Intratracheal inoculation of adenoviral-Cre (AdCre) was used to simultaneously activate the conditional Kras allele (*LSL-KRAS^G12D^*) and delete *p53* (*p53^fl/fl^*) *in vivo* and induce lung tumorigenesis (49). The *Rosa26:LSL-MAGEA4-GFP* mouse line was designed and generated at the UNC Animal Models Core Facility. Schematic diagrams were generated using BioRender (https://www.biorender.com).

### Genotoxin administration

Mice received 50 ug/ml 4-Nitroquinoline N-oxide (4NQO, Sigma N8141) dissolved in 20 μg/ml 1,2 propanediol (propylene glycol, Sigma 398039) diluted with sterile water. A roughly 150 to 250 mL of 4NQO solution was given for every animal cage, with the amount varying based on the quantity and weight of animals within each cage. Mice were humanely euthanized under the following circumstances: 1) if they had excessively large tumors >2cm outside the oral cavity or smaller tumors that hindered their ability to consume food or water, 2) in cases of significant weight loss exceeding 20% of the animal’s initial weight before the study, or 3) when an observer determined the presence of a moribund state, characterized by signs such as lethargy, a hunched posture, and/or indications of pain/wincing.

### Tissue Collection and Processing

For collection of relevant tissues for downstream processes and analysis, mice were humanely euthanized in accordance with IACUC and UNC Division of Comparative Medicine (DCM) guidelines.

### RNA-sequencing (RNA-seq) analysis

RNA was extracted from the cells isolated from mouse tongue epithelium using either the Qiagen RNeasy or AllPrep kit, according to the manufacturer’s instructions. Library preparation and sequencing was performed by Genewiz. Raw sequencing reads were trimmed and aligned to the Mus musculus reference genome (GRCm38/mm10 assembly) using the BBMap package (v37.00). Resulting SAM files were converted to BAM format and sorted using SAMtools v1.3.1. Read quantification and matrix generation were carried out with featureCounts. Count matrices for weeks 10, 16, 22 and 29 were obtained from a previously published transcriptional profiling study by *Lee et al.* (39). Differential gene expression analysis was performed using DESeq2, and functional enrichment of significantly altered gene sets was assessed using PANTHER. The TCGA Head and Neck Squamous Cell Carcinoma (HNSC) RNA-Seq data and clinical data for primary tumor and solid tissue normal were accessed using R (version 4.3.3) and the TCGAbiolinks package (version 2.30.0). TCGA variant data was obtained from cBioPortal. DNA damage response genes and pathways used in the study are provided as Supplementary Table 2.

Heatmaps from RNAseq data were generated from the log transformed and normalized gene expression matrix using ComplexHeatmap package (version 2.18.0) in R (v4.3.1).

### Whole exome sequencing (WES)

For 4NQO-treated mice, DNA was extracted from tumors utilizing the Zymo Quick-DNA Microprep Plus Kit (D4074, Zymo) according to the manufacturer’s guidelines. Isolated DNA samples were submitted to Genewiz or UNC High-Throughput Sequencing Facility for library preparation and sequencing. For KRAS-induced lung tumors, DNA was extracted from tumors using the DNeasy Blood and Tissue Kit (69506, Qiagen). The DNA samples were sent to Novogene for library preparation and sequencing. Analysis of exome sequencing data and quantification of SNVs and INDELs was performed using the computational pipeline developed by Lange et al. For bioinformatic analysis, initial quality check of raw sequencing reads (FASTQ files) was performed using FastQC(version 0.11.8) (https://www.bioinformatics.babraham.ac.uk/projects/fastqc/). Adapter sequences and low-quality bases were removed using Trimmomatic (version 0.39) (50). FastQC was re-evaluated on the trimmed reads for quality check. Trimmed reads were then aligned to mouse reference genome, GRCm38 (VM25), using BWA-MEM (version 0.7.17) (51). The resulting sequence alignment map (SAM) files were converted to binary alignment map (BAM) files, sorted, and cleaned using Picard (version 2.21.7) (http://broadinstitute.github.io/picard). Read group information was added, and duplicate reads were marked without removal. To detect systematic errors made by the sequencing machine and correct the base quality score, Base Quality Score Recalibration (BQSR) was performed using BaseRecalibrator and ApplyBQSR tools within GATK (version 4.1.7.0) (52), based on known Single Nucleotide Polymorphism (SNP) and small insertions/deletions (indel) sites from the Mouse Genomes Project (REL-1505) specific to the C57BL/6NJ mouse strain. Quality control of recalibrated BAM files was performed using Picard’s CollectSequencingArtifactMetrics, CollectMultipleMetrics, BedToIntervalList, and CollectHsMetrics functions as well as SAMtool’ (version 1.9a) (53) idxstats function. Single nucleotide variants (SNVs) and indels in the samples were identified using Mutect2 tool of GATK. Probable germline artifacts, FFPR artifacts (C/T), and oxidative DNA damage artifacts (G/T) were filtered using FilterMutectCalls function, FilterByOrientationBias function, and SnpSift (v4.3t). Variants with indel length >10 base pairs were excluded using GATK SelectVariants. Additional filtering was applied using SnpSift (v4.3t), retaining variants with a tumor allele frequency (AF) ≥ 10%, a minimum read depth of 10, and at least 3 supporting alternate reads. Analysis and visualization of mutational signatures from VCF files was performed using MutaionalPatterns (v3.12.0) (54) in R (v4.3.1) or SomaticSignatures (v2.40.0)(55) in R (v4.4.0).

## RESULTS

### RAD18 confers 4NQO-tolerance in oral keratinocytes

Chronic (8-16 week) treatment with 4NQO is widely used to induce oral squamous cell carcinomas (OSCCs) in experimental mice.(38,56–60) We evaluated the 4NQO-induced OSCC model as an experimental system for defining Rad18 roles in carcinogenesis. First, we tested the potential role of RAD18 in mediating responses to 4NQO-induced DNA damage in two different human oral keratinocytes, OKF-6 (normal) and DOK (dysplastic). 4NQO-treated human oral keratinocyte cells showed an increase in PCNA mono-ubiquitination when compared with control cultures (Fig. 1A). Importantly, 4NQO-induced PCNA mono-ubiquitination was attenuated in RAD18-depleted cells, showing that RAD18 mediates 4NQO-induced PCNA mono-ubiquitination. In RAD18-depleted cells, 4NQO-treatment led to an increase in activation-associated phosphorylation of the DNA Double-Stranded Break (DSB)-sensing protein kinase ATM. In OKF-6 cells, the combination of 4NQO and RAD18 knockdown led to elevated phosphorylation of γH2AX and CHK1. In the dysplastic DOK cells (which carry a *TP53* mutation), yH2AX levels were basally high and not further induced by 4NQO and RAD18-loss.

**Figure 1.**
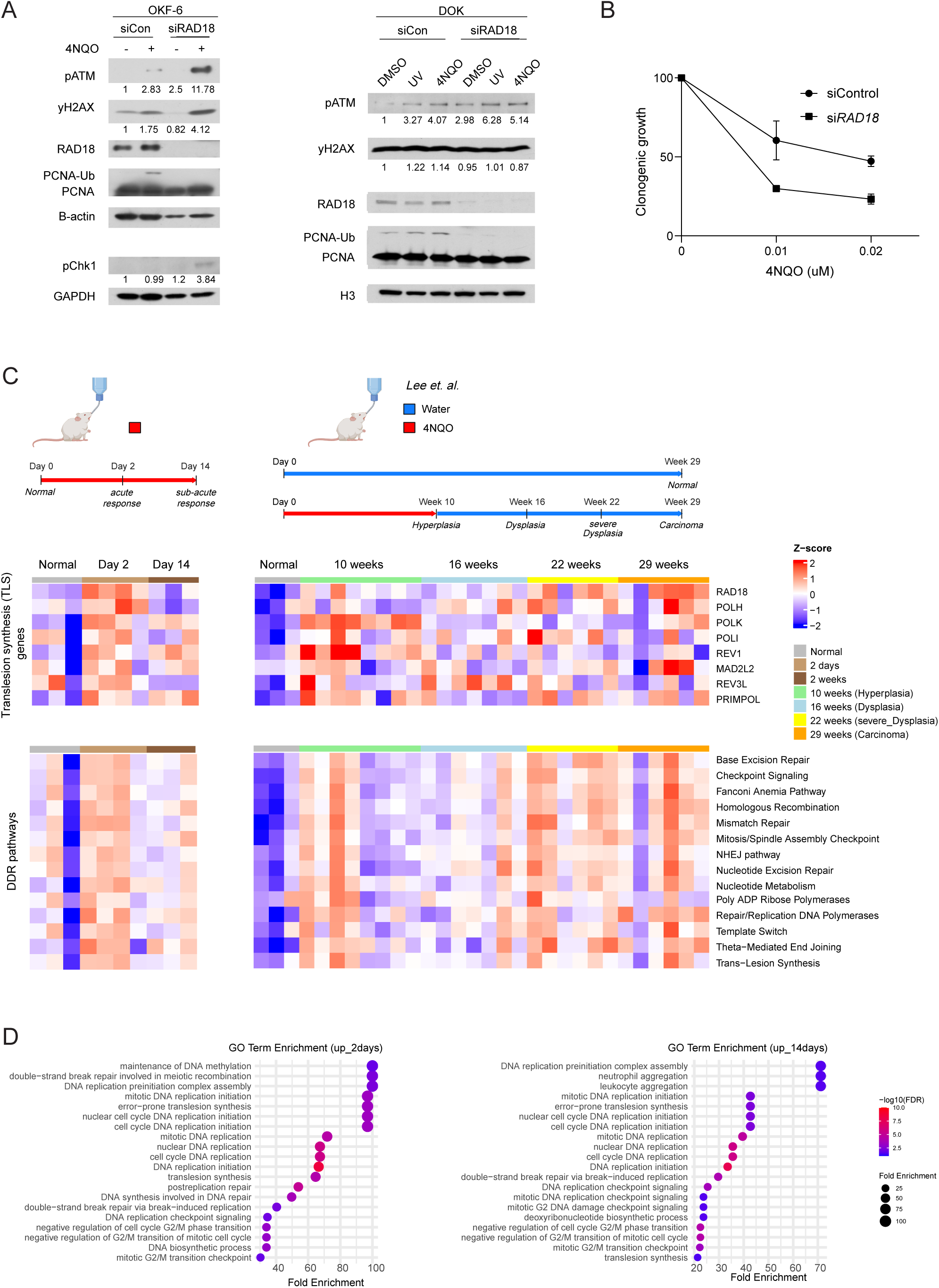
Rad18 promotes tolerance of 4NQO-nduced DNA damage *in vitro* and is elevated during 4NQO-induced oral carcinogenesis. (A) OKF-6 (normal oral keratinocytes) and DOK (dysplastic oral keratinocytes) cells were transfected with control non-targeting siRNA (siCon) or RAD18-directed siRNA, then treated with UV (20 J/m^2^) or 4NQO (0.01μM). 4 h after genotoxin treatment, cell lysates were prepared and analyzed by SDS-PAGE and immunoblotting. (B) DOK cells were treated with siCon or siRAD18, then tested for 4NQO-sensitivity using clonogenic survival assays. Error bars represent the standard deviation (SD) of the mean. Statistical analysis: two-way ANOVA followed by Sidak’s multiple comparison test, \**P* < .05, \*\**P* < .01, \*\*\**P* < .001, \*\*\*\**P* < .0001, ns = not significant. (C) Scheme showing *in vivo* 4NQO treatment regimens in the current study (50 ug/ml 4NQO) and study by *Lee et al*. (100 ug/ml 4NQO). In our study, *Rad18^+/+^*mice were treated with 4NQO for 2 days or 14 days (or left untreated for controls). Then tongue epithelia from all three treatment groups were subjected to transcriptional profiling. Heatmap of TLS and DDR pathway genes in the 4NQO treatment groups when compared with control mice in our study and study by *Lee et al*. (D) Gene ontology (GO) analysis of upregulated genes at day 2 and 14 in 4NQO-treated oral epithelium showing top 20 GO terms.

Overall, the results of Fig. 1A suggest that RAD18-mediated TLS averts ‘collapse’ of DNA replication forks that encounter 4NQO-induced DNA damage into, as we have previously shown for other replication fork-stalling DNA lesions (45,61). In clonogenic survival assays, RAD18-depletion led to 4NQO-sensitivity (Fig. 1B), demonstrating that RAD18 confers tolerance of 4NQO-induced genotoxicity. Therefore, RAD18 is an important component of the DNA damage response to 4NQO in oral keratinocytes.

### Expression of Rad18 and TLS pathway genes is dynamically regulated in vivo in response to 4NQO treatment

Our analysis of TCGA patient data revealed that HPV-negative human head and neck squamous cell carcinomas (HNSCs) show a small but statistically significant increase in *RAD18* expression relative to normal adjacent tissue (Supplementary Fig. S1). To test the potential involvement of *Rad18* and other TLS pathway genes in 4NQO-induced OSCC development we performed transcriptional profiling of basal keratinocytes isolated from the tongues of mice that were treated for 0 days, 2 days, or 14 days with 4NQO (Fig. 1C, upper scheme). The first (2 day) timepoint of 4NQO exposure was chosen to allow sufficient time for the animals to drink the 4NQO-laden water. The second timepoint of 14 days of exposure was chosen to allow for tissue-level changes and homeostasis to be reestablished in the presence of the genotoxin, prior to the emergence of malignant lesions. We performed bulk RNA-sequencing (RNA-seq), then evaluated the differentially expressed genes at each 4NQO treatment timepoint using DESeq2 (62).

After 2 days of 4NQO exposure we identified 145 transcripts that were downregulated and 131 mRNAs that were upregulated (log2 fold change >|1|, false discovery rate p<0.05) in 4NQO-exposed oral keratinocytes when compared with control untreated mice. After 14 days of 4NQO exposure, 317 genes were upregulated, and 283 genes were downregulated in oral keratinocytes from 4NQO-treated mice when compared with the control (untreated) group. When we plotted the gene expression data based on their principal component analysis (PCA), the three treatment groups (baseline, 2 days, and 14 days 4NQO) clustered very distinctly (Supplementary Fig. 2A), suggesting that the 4NQO-induced transcriptional program is temporally controlled. We analyzed the differentially-expressed gene lists for statistical overrepresentation using PANTHER (63,64) to identify the biological pathways that were altered by 4NQO treatment (Fig. 1D, Supplementary Table 1) (Scott). Interestingly, in samples from the 2-day 4NQO treatment group, TLS was one of the most over-represented biological processes (FDR p=2.28 x 10^-2^), based on overexpression of *Rad18* (padj = 0.014), and the Y-family TLS polymerases *Polk* (padj = 8.93×10^-5^) and *Rev1* (padj = 0.003). Moreover, *Polk* and *Rev1* mRNAs remained upregulated after 14 days of 4NQO treatment (padj = 2.9 x 10^-4^ and padj = 1.49 x 10^-5^, respectively). In addition to TLS, GO terms related to DNA replication, cell cycle and DSB repair were enriched in 4NQO-treated oral epithelium, suggesting that 4NQO genotoxicity induces a coordinated transcriptional program of DDR genes to promote genome maintenance (Fig. 1D). Unique to the 14-day timepoint were GO terms related to immune infiltration such as neutrophil and leukocyte aggregation (Supplementary Fig. 2B). The heatmap in Fig. 1C (upper left panel) shows the relative changes in the expression of individual TLS pathway genes (including *Rad18*, *Polh*, *Polk*, *Poli*, *Rev1*, *Mad2l2*, *Rev3l* and *Primpol*) at 2 days and 2 weeks post-4NQO treatment. We also used previously-curated gene panels (65) to interrogate expression of TLS and other DDR transcripts at the pathway level. As shown in Fig. 1C (lower left panel) and Supplementary Fig. 3, acute (2d) 4NQO treatment led to increased expression of mRNAs encoding core components of all major genome maintenance pathways.

To complement our *in vivo* analysis of oral keratinocytes that were treated with 4NQO acutely (2 weeks, corresponding to early pre-neoplasia), we analyzed TLS/DDR transcriptomes using an independent RNASeq dataset(39) derived from mice that were subject to a chronic (up to 10 weeks) 4NQO treatment and then allowed to develop oral tumors (Fig. 1C, lower scheme).

As shown in Fig. 1C (right side panels) and Supplementary Fig. 4, mRNA levels of DDR genes (including TLS genes) were broadly elevated in most/more than half of the samples at week 10 (a time at which 4NQO treatment was stopped and hyperplasia had developed). At week 16 (six weeks after 4NQO treatment was terminated and dysplasia was evident), the expression of DDR pathway mRNAs declined to near normal levels, potentially reflecting the reestablishment of homeostasis in the absence of chemically-induced genotoxicity. Interestingly, at weeks 22 and 29 (times associated with severe dysplasia and carcinoma respectively), expression levels of DDR genes were once again elevated (Fig. 1C). The increased expression of DDR genes in the severely dysplastic (week 22) and malignant (week 29) lesions may reflect a greater demand for genome maintenance during the later stages of cancer to promote tolerance of intrinsic replication stresses.

Taken together, the results of Fig. 1C show that genes encoding TLS and other DDR factors are dynamically upregulated *in vivo* following 4NQO-induced DNA damage and during dysplasia and malignancy. Notably, the elevated expression of *Rad18* that occurs in a mouse model of chemically-induced OSCC recapitulates the increased expression of *RAD18* observed in human HPV-negative oral cancers.

### Rad18 suppresses 4NQO-induced oral carcinogenesis in vivo

Our findings show that RAD18 increase survival of 4NQO-treated cells (Fig. 1A, 1B), and that *Rad18* and other TLS pathway mRNAs are dynamically regulated in response to carcinogen treatment (Fig. 1C) suggested a potential role for Rad18 in modulating 4NQO-induced oral carcinogenesis. To test the role of Rad18 in 4NQO-induced oral carcinogenesis we performed tumorigenesis studies comparing cohorts of *Rad18^+/+^* control mice with *Rad18^-/-^* KO and *Rad18^+/-^* heterogenous mice. The design of our carcinogenesis studies is illustrated in Fig. 2A, with 8 weeks of 4NQO exposure in drinking water followed by regular monitoring. Interestingly, tumorigenesis was significantly more rapid in *Rad18^-/-^* mice when compared with *Rad18^+/^* (p = 0.006) (Fig. 2B), *Rad18^+/+^* (p = 0.0338) (Supplementary Fig. 5A) and *Rad18^+/-^*(p = 0.0021) mice (Supplementary Fig. 5B). Immunostaining experiments showed a trend of elevated γH2AX levels in *Rad18^-/-^* tumors when compared with *Rad18^+/+^* tumors and the adjacent tongue tissue from *Rad18^-/-^* mice (Fig. 2C). However, the difference in γH2AX staining between genotypes did not reach statistical significance. Nevertheless, these data show that *Rad18* suppresses 4NQO-induced oral carcinogenesis.

**Figure 2.**
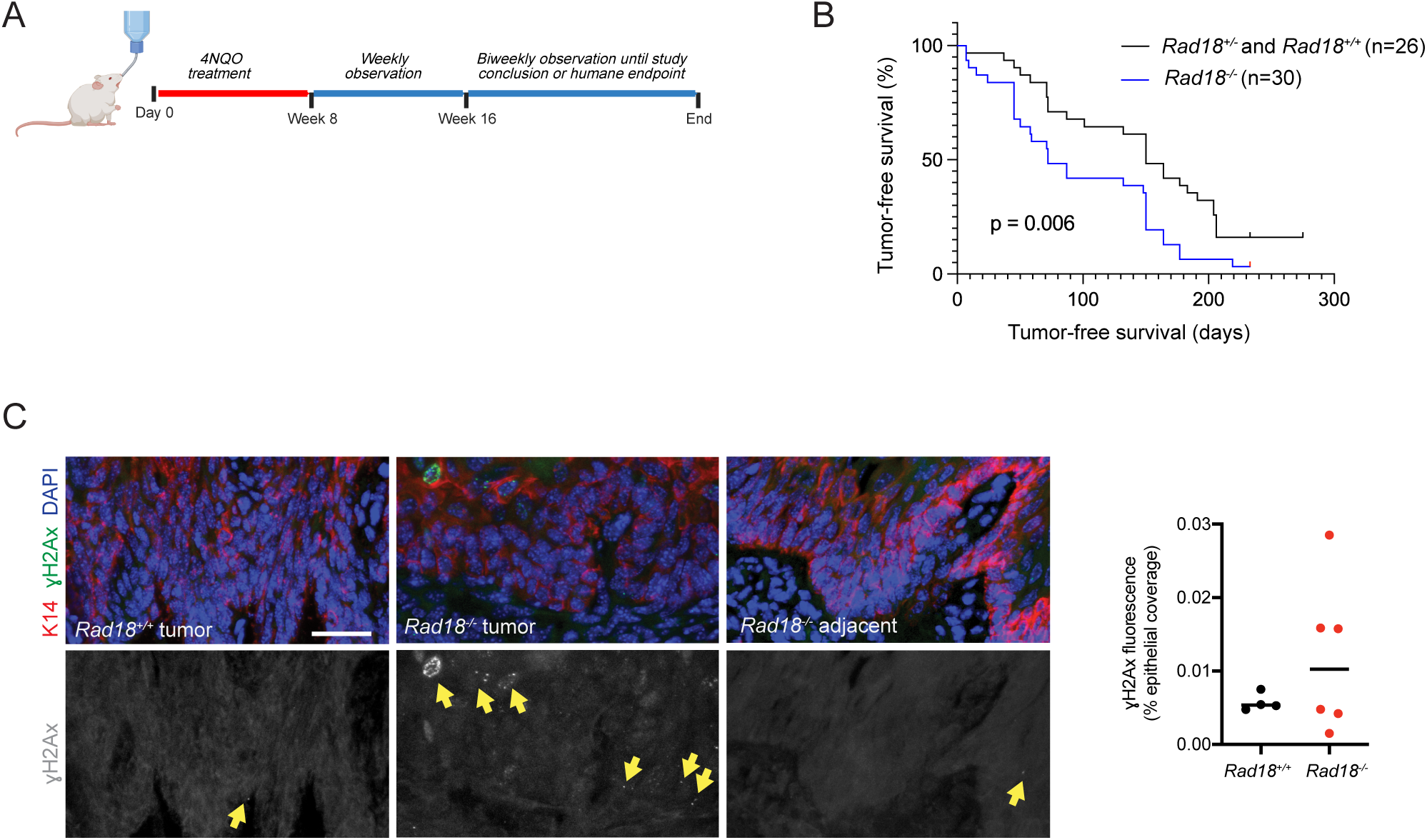
*Rad18* suppresses 4NQO-induced oral carcinogenesis. (A) Scheme showing *in vivo* 4NQO treatment regimen in our study. (B) Survival curve showing that *Rad18^-/-^*mice develop tumors more rapidly than Rad18-expressing *Rad18^+/^* littermates (*p* = 0.006). The *Rad18^+/^* cohort comprises *Rad18^+/+^* and *Rad18^+/-^* animals. Log-rank (Mantel-Cox) tests were performed to assess the statistical significance of survival curves. (C) Immunostaining experiment showing a trend of elevated γH2AX levels in *Rad18^-/-^* tumors when compared with *Rad18^+/+^* tumors and the adjacent tongue tissue from *Rad18^-/-^* mice.

### Rad18 represses 4NQO-induced G(C)>T(A) transversions

Tumor-suppressive genome maintenance proteins typically promote error-free DNA repair and avert formation of cancer-promoting mutations. To determine the effect of *Rad18* on mutagenesis we sequenced exomes of 4NQO-induced oral tumors from *Rad18^+/+^*(n=3), *Rad18^+/-^* (n=4), and *Rad18^-/-^* (n=7) mice. We used matched-normal tail epithelium from each animal to generate a reference genome and deployed a previously described computational pipeline (36) to annotate and quantify mutations.

First, we assessed the number and types of single nucleotide variation (SNV) in the genotype groups. *Rad18^-/-^*mice had the highest total SNV burden (5820 +/- 1555) and WT mice had the lowest (3642 +/- 1389), with *Rad18^+/-^* demonstrating an intermediate phenotype. These differences, however, did not achieve statistical significance by ANOVA (p=0.0662 for *Rad18^-/-^* vs *Rad18^+/+^*) (Fig. 3A). As expected, G(C)>T(A) transversions (known 4NQO-inducible SNVs) represented the most abundant SNV in oral cancers from all genotypes. G(C)>T(A) transversions were significantly more abundant in *Rad18^-/-^* tumors than either *Rad18^+/-^* or *Rad18^+/+^* tumors (Fig. 3B; *Rad18^-/-^* vs *Rad18^+/-^*, padj = 0.04; *Rad18^+/-^* vs *Rad18^+/+^*, padj = 0.007; *Rad18^-/-^* vs *Rad18^+/+^*, padj < 0.0001). In contrast with SNVs, frequencies of various insertion/deletion (indel) mutations in 4NQO-induced tumors were not significantly affected by Rad18 (Fig. 3C). Interestingly, *Rad18^-/-^*mice had the highest average DBS burden (43.25 +/- 14.2) and WT mice had the lowest (23.66 +/- 1389), with *Rad18^+/-^* having an intermediate burden (33.33 +/- 7.23) (Fig. 3D). These differences, however, did not achieve statistical significance by ANOVA (p=0.0916 for *Rad18^-/-^* vs *Rad18^+/+^*) (Fig. 3D). CC>NN transversions were most abundant across all the tumors and were significantly more abundant in *Rad18^-/-^*tumors than *Rad18^+/+^* tumors (Fig. 3D; *Rad18^-/-^* vs *Rad18^+/-^*, p = 0.0255), particularly for CC>AA transversions which are associated with cosmic double base substitution (DBS) signature DBS2 (etiology: tobacco smoking and other mutagen) (Fig. 3D). Next, we determined the mutational signatures of 4NQO-induced tumors based on signatures curated in the COSMIC v3 database. As shown in Fig. 3E, Single Base Substitution signature 4 (SBS4, which is associated with tobacco smoking in HNSC), was the predominant 4NQO-induced mutational signature across all samples. SBS29, which is associated with tobacco chewing, was the second largest contributor to the overall mutational signatures of all samples (Fig. 3E). We did not observe any statistically significant difference in the patterns of mutational signatures between different genotypes (Fig. 3E). Taken together, the results of Fig. 3 suggest that Rad18 promotes error-free replication and repair of 4NQO-induced lesions, likely explaining why Rad18 suppresses 4NQO-induced carcinogenesis.

**Figure 3.**
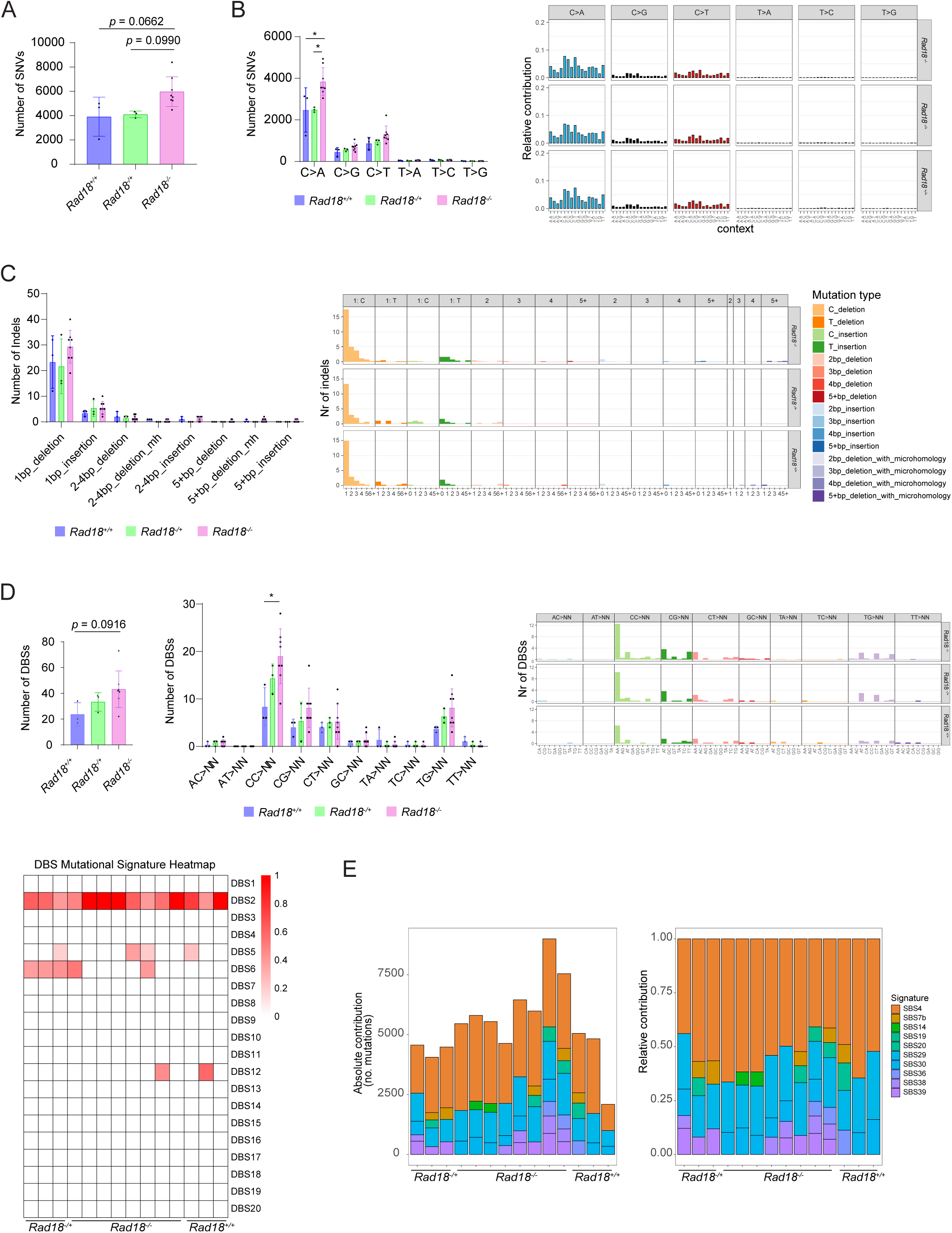
*Rad18* loss leads to altered mutation patterns mouse OSCCs. (A) WES analysis of single nucleotide variants (SNV) in mouse oral tumors harvested from *Rad18^+/-^* (het), *Rad18^-/-^* (KO), and *Rad18^+/+^* (WT) animals. The data show that *Rad18^-/-^*tumors had more 4NQO-mediated mutations compared to *Rad18^+/+^ (p* = 0.0662). Error bars represent the standard deviation (SD) of the mean. Statistical analysis was performed using ANOVA with Tukey’s secondary test. Statistical analysis using t test between *Rad18^-/-^* (KO), and *Rad18^+/+^*(WT) showed statistical significance (*p* = 0.0467). (B) The data show that *Rad18^-/-^* tumors had more G(C)>T(A) transversions than *Rad18^+/-^* (padj = 0.04) or *Rad18^+/+^* (*p* < 0.001). *Rad18^+/-^* tumors also had significantly more G(C)>T(A) transversions than *Rad18^+/+^*tumors *(p* = 0.007). Error bars represent the standard deviation (SD) of the mean. Statistical analysis was performed using ANOVA with Tukey’s secondary test. (C) Statistical analysis of insertions and deletions (INDELs) in mouse oral tumors from the same samples and genotype groups as in (A). No statistically- significant differences were evident when comparing indel numbers between experimental groups. (D) The data show that *Rad18^-/-^* tumors had more double base substitutions (DBSs) than *Rad18^+/+^* (*p* = 0.0916). Error bars represent the standard deviation (SD) of the mean. Statistical analysis was performed using ANOVA with Tukey’s secondary test. Statistical analysis using t test between *Rad18^-/-^*(KO), and *Rad18^+/+^* (WT) showed *p* = 0.0566. *Rad18^-/-^*tumors also had significantly more CC>NN transversions than *Rad18^+/+^*tumors (*p* = 0.007). Statistical analysis was performed using ANOVA with Tukey’s secondary test. (E) Stacked bar chart showing the absolute and relative contribution of COSMIC SBS mutational signatures to the mutational profile of 4NQO-induced oral tumors collected from *Rad18^+/-^* (het), *Rad18^-/-^* (KO), and *Rad18^+/+^*WT animals.

### The pathological Rad18 activator MAGEA4 suppresses 4NQO-induced carcinogenesis

Many human cancer cells aberrantly express high levels of the germ cell protein Melanoma Antigen A4 (MAGE-A4) which is a binding partner and activator of RAD18 (35). Many cancer cells rely on MAGE-A4 to induce high-level RAD18 expression and sustain TLS (35). Our analysis of TCGA patient data revealed that HPV-negative human head and neck squamous cell carcinoma (HNSCC) tumors show increased expression of type I MAGEs (which comprise MAGE-A, -B, and -C subfamilies) when compared with normal adjacent tissue. Intriguingly, *MAGEA4* was one of the top 50 genes differentially overexpressed in HNSC HPV negative tumors (Fig. 4A). Therefore, *MAGEA4* expression is a feature of many oral cancer cells. These results may indicate that HPV-positive tumors adapt to oncogenic stress by increasing *RAD18* mRNA expression, whereas HPV-negative tumors rely on MAGEA4 for stabilizing and activating RAD18 at the protein level.

**Figure 4.**
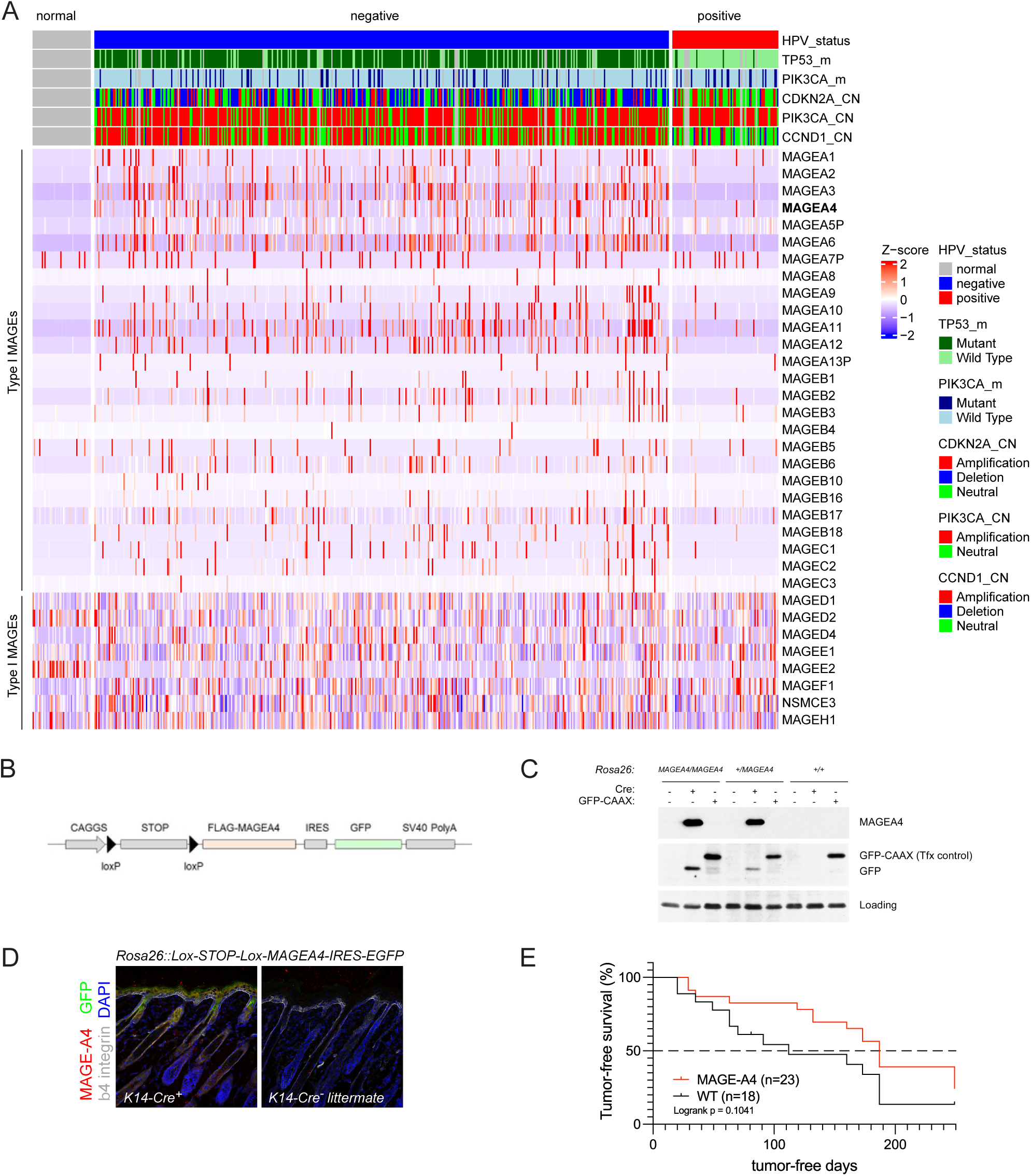
Development of a transgenic mouse model for conditional overexpression of MAGEA4. (A) Heatmap of mRNA expression of Type I and Type II MAGE genes in the TCGA-HNSC HPV negative and positive tumors when compared with adjacent normal control. (B) Design of targeting vector used to drive Cre-inducible MAGEA4 expression from the *Rosa26* locus. (C) Validation of MAGEA4 and GFP expression in *Rosa26^+/LSL-MAGEA4^* and *Rosa26^LSL-MAGEA4/LSL-MAGEA4^* MEF after transient transfection with a CMV-Cre expression vector. GFP-CAAX was used as a transfection control. (D) Validation of Cre-inducible MAGEA4 and GFP expression in skin sections from *K14-Cre; Rosa26^+/LSL-MAGEA4^* mice (left panel). The right panel is a control experiment showing absence of GFP and MAGEA4 immunostaining in a skin section from a *Rosa26^+/LSL-MAGEA4^* littermate not expressing Krt14-Cre. (E) Survival curve showing rates of 4NQO-induced oral carcinogenesis in *K14-Cre; Rosa26^+/LSL-MAGEA4^* mice and control *Rosa26^+/LSL-MAGEA4^* littermates. Log-rank (Mantel-Cox) test was performed to assess the statistical significance of survival curves.

To complement our loss-of-function experiments (in which we defined the effects of *Rad18*-deficiency), we devised a gain-of-function strategy to test how Rad18 activation by MAGE-A4 *in vivo* impacts 4NQO-induced mutagenesis and oral carcinogenesis. We generated a transgenic vector for Cre-inducible expression of MAGE-A4 (and a co-expressed GFP allele on a polycistronic transcript) from the *Rosa26* locus *in vivo* (Fig. 4B). We validated Cre-inducible expression of MAGE-A4 (and GFP) in mouse embryonic fibroblasts (MEF) from transgenic mice (Fig. 4C). Ectopically-expressed MAGEA4 and GFP were also readily detectable in skin sections from Keratin 14-Cre-expressing (Krt14-Cre) *Rosa26^+/LSL-MAGEA4^* transgenic mice (Fig. 4D).

Having validated Cre-dependent expression of the MAGEA4 transgene, we performed 4NQO-tumorigenesis studies using cohorts of *Krt14-Cre*; *Rosa26^+/LSL-MAGEA4^* mice (n=23) and strain-matched *Rosa26^+/LSL-MAGEA4^*control animals lacking the *Krt14-Cre* transgene (n=18). As shown in Fig. 4E, rates of 4NQO-induced tumorigenesis were lower in MAGEA4-expressing (*Krt14-Cre*; *Rosa26^+/LSL-MAGEA4^)* mice when compared with the control *Rosa26^+/LSL-MAGEA4^* cohort (*p* =0.1041). Interestingly, when only male mice were considered, the survival difference was significant (*p* =0.007; median survival of 35.5 months for *Krt14-Cre*; *Rosa26^+/LSL-MAGEA4^* males compared to 119 for control males), indicating a gender-specific effect of MAGE-A4 expression on tumorigenesis in this model system.

We harvested oral tumors from the control and MAGEA4-overexpressing mice and sequenced their exomes. We used normal tail epithelium from every tumor-bearing mouse to generate a matched reference genome and annotated mutations in all tumors. The total and types of SNVs were not significantly different between wild-type and MAGEA4-overexpressing tumors (Fig. 5A and 5B). Our analysis of insertions and deletions (INDELs) indicated a trend toward fewer INDELs in *Krt14-Cre*; *Rosa26^+/LSL-MAGEA4^* tumors compared to *Rosa26^+/LSL-MAGEA4^*, but this trend only reached significance for 2-4bp_insertions (*p* = 0.0256) though the number of mutations were low (*Rosa26^+/LSL-MAGEA4^*1.00 ± 0.71; *Krt14-Cre*; *Rosa26^+/LSL-MAGEA4^* 0.125 ± 0.35) (Fig. 5C). As expected, SBS4 and SBS29 were the predominant mutational signatures in all 4NQO-induced tumors. However, there were no statistically significant differences in the relative abundance of different mutational signatures between *Krt14-Cre*; *Rosa26^+/LSL-MAGEA4^* and *Rosa26^+/LSL-MAGEA4^* tumors (Fig. 5D). Taken together, the result of our tumorigenesis studies with MAGEA4-overexpressing mice is consistent with previous data from Fig. 3 however, MAGEA4 overexpression did not reduce 4NQO-induced mutations.

**Figure 5.**
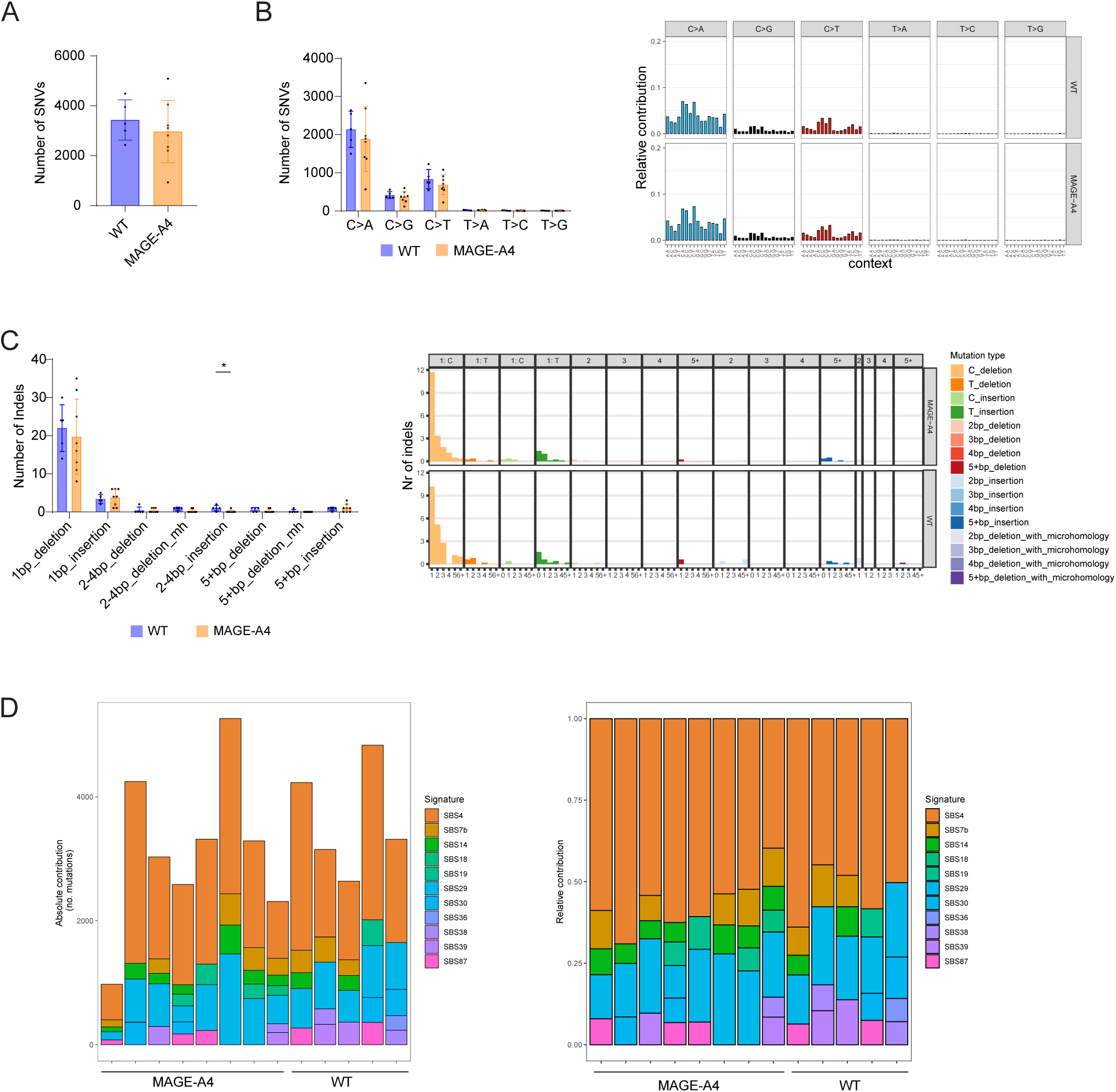
Effect of MAGEA4 overexpression on mutational spectra of 4NQO-induced oral cancers. (A-B) WES analysis of oral tumors collected from *K14-Cre; Rosa26^+/LSL-MAGEA4^* mice and control *Rosa26^+/LSL-MAGEA4^* littermates showing similar levels of mutations and a predominance of C>A and C>T transversions in both genotypes. Error bars represent the standard deviation (SD) of the mean. Statistical analysis was performed using Mann-Whitney test. (C) Statistical analysis of insertions and deletions (INDELs) in mouse oral tumors from the same samples and genotype groups as in (A). Error bars represent the standard deviation (SD) of the mean. Statistical analysis was performed using Mann-Whitney test. No statistically- significant differences were evident when comparing indel numbers between experimental groups. (D) Stacked bar chart showing the absolute and relative contribution of COSMIC SBS mutational signatures to the mutational profile of 4NQO-induced oral tumors collected from *K14-Cre; Rosa26^+/LSL-MAGEA4^* mice and control *Rosa26^+/LSL-MAGEA4^*littermates.

### Effect of Rad18 on genomic landscape of Kras-induced lung tumors

For the purpose of comparison with our mutagen (4NQO)-induced carcinogenesis studies, we next tested the impact of Rad18 in an oncogene-induced tumorigenesis model. We opted to study the potential role of Rad18 in Kras-induced lung tumorigenesis as RAD18 is typically overexpressed in lung adenocarcinomas and positively correlated with SNVs irrespective of smoking history (22). Moreover, we have previously shown that Rad18 is important for averting single-stranded DNA (ssDNA) formation and sustaining proliferation of oncogenic Kras-expressing mouse embryonic fibroblasts (33). Therefore, we tested our hypothesis that Rad18 would facilitate Kras-induced lung tumorigenesis. We used adenovirus encoding Cre to activate a latent *KRas^G12D^*allele in the lungs of *Rad18^+/+^*; *p53^-/-^* (n = 7) and *Rad18^-/-^*; *p53^-/-^* (n = 24) mice, then measured rates of tumorigenesis in the two cohorts. We performed these experiments using a p53-deficient background to model the genetics of lung cancer in humans, and because we have previously shown that abrogation of the TP53-mediated G1 checkpoint creates a dependency on RAD18 for DNA damage tolerance (33). Unexpectedly, *Rad18* status did not significantly affect survival rates (based on humane endpoint criteria) of mice harboring *Kras^G12D^*-induced lung tumors (Supplementary Fig. 6A). To determine whether Rad18 affected genomic characteristics of *Kras^G12D^*-induced tumors, we sequenced lung lesions from *Rad18^+/+^* and *Rad18^-/-^*mice and computationally annotated SNVs and Indels in tumor exomes from the two cohorts. As shown in Supplementary Fig. 6C, G(C)>A(T) substitutions were the most abundant substitutions in *Kras^G12D^*-driven lung tumors, regardless of Rad18 status. However, neither the total SNV burdens (Supplementary Fig. 6B), nor the frequencies of any individual mutations (Supplementary Fig. 6C) were significantly different between *Rad18^+/+^*and *Rad18^-/-^* tumor exomes. Notably, the G(C)>T(A) transversions that were repressed by Rad18 in the 4NQO-induced oral cancers (Fig.5B), were equally abundant in *Rad18^+/+^* and *Rad18^-/-^* lung tumors that were *Kras^G12D^*-driven (Supplementary Fig. 6C). Therefore, Rad18 can have different effects on frequency of a specific SNV in tumors depending on the nature of the cancer-inciting agent and / or tissue context. We also compared the indel spectra between *Kras^G12D^*-induced *Rad18^+/+^* and *Rad18^-/-^* lung tumors. As shown in Supplementary Fig. 6D, only 1bp insertions (but not other indels) were higher in *Rad18^-/-^* lung tumors when compared with *Rad18^+/+^*. Therefore, in *Kras^G12D^*-driven tumors, Rad18 promotes error-free DNA replication and/or repair events to avert insertion mutations.

## DISCUSSION

It is well established that rapid changes in DDR signaling (i.e. occurring within minutes of DNA damage) are mediated via dynamic post-translational modifications of DNA damage sensing and transducing proteins (66–68). On the other hand, more gradual and long-term adaptive reprogramming of genome maintenance might be achieved via changes in the expression levels of core DDR factors in response to altered demands of cells. Here using a well-established 4NQO-induced oral carcinogenesis model we define a biphasic pattern of DDR gene induction that is evident both early during the genotoxin-treatment phase, and as a discrete distal wave that coincides with dysplasia and malignancy. We have previously shown that cancers can be subtyped based on their DDR transcriptomes which in turn reflect the unique DNA damaging exposures and intrinsic DNAreplication stresses associated with each malignancy (65,69). Our temporal analysis of the DDR transcriptome reveals how the reprogrammed genome maintenance networks of malignant cells evolve during multi-step chemically-induced oral carcinogenesis. Interestingly, we show here that transcriptional induction of *Rad18* is associated with both waves of DDR gene induction and persists in the established malignant 4NQO-induced OSCC. High level expression of *RAD18* and of its pathological activator *MAGEA4* are also features of many human cancers (22) including HNSCC as shown here.

RAD18-mediated TLS has the potential to promote two major enabling characteristics of cancer cells: (i) mutagenesis (which occurs via error-prone replication and repair of carcinogen-damaged DNA) and (ii) tolerance of oncogene-induced DNA replication stress (which results from remediation of oncogene-induced replication fork stalling). The temporal patterns of *Rad18* expression that we observe during carcinogenesis in this model are consistent with potential roles for Rad18 in responding to both 4NQO-induced genotoxicity during pre-neoplasia as well as intrinsic DNA replication stress that arises later in transformed cells.

Based on its roles in promoting mutagenesis and tolerance of oncogenic stress it is reasonable to hypothesize that Rad18 might stimulate carcinogenesis. However, the limited number of studies done to address the role of *Rad18 in vivo* have not yielded a singular paradigm for how Rad18 affects mutagenesis and tumorigenesis. In a DMBA-induced skin carcinogenesis model, Rad18-deficiency led to reduced levels of A(T)>T(A) SNVs while increasing the formation of small 4bp deletions. Thus, for DMBA-induced DNA damage, Rad18 promotes error-prone (mutagenic) DNA replication that induces SNVs while averting Indels (36).

In contrast, in the 4NQO-induced oral tumorigenesis model described here, *Rad18* loss led to increased numbers of C>A transversions (and increased rates of tumorigenesis). Therefore, Rad18 promotes error-free replication of 4NQO-induced DNA lesions thereby averting mutations. Our observation that the Rad18 activator MAGE-A4 (35) also reduces 4NQO-induced carcinogenesis but do not affect rates of 4NQO-induced mutagenesis is interesting and points to a nuanced role of MAGEA4 requiring further analysis. Interestingly, in the context of DNA damage induced by the chemotherapy agent temozolomide (TMZ) in glioblastoma cells, RAD18 promotes error-free bypass of O6mG while simultaneously allowing error-prone bypass of other TMZ-induced lesions (N^7^mG and N^3^mA) (29). Collectively these observations suggest that the impact of Rad18 on error-propensity of DNA replication (and on tumorigenesis) is highly context-dependent and determined by the nature of the DNA damage.

4NQO is known to generate tobacco related cosmic signature 4 (38,39) and indeed in our experiment, cosmic signature 4 was the dominant signature in both *Rad18^+/+^* and *Rad18^-/-^*backgrounds. Therefore, Rad18 does not specify the hallmark 4NQO-induced mutational signature. Since G>T transversions are enriched in Rad18-deficient mice, it is surprising that the contribution of SBS4 (a signature defined by G to T mutations) is unaffected by *Rad18* status. We must conclude that SBS4 is mediated via an alternative Rad18-independent mutagenic mechanism. We hypothesize that at least three putative mechanisms could explain the Rad18-independence of SBS4: First, alternative PCNA-directed E3 ubiquitin ligases such as HLTF and CRL4/Cdt2 might promote PCNA ubiquitylation and mutagenesis via the Y-family TLS polymerases in cells lacking Rad18 (70,71). Second, although Rad18 promotes the recruitment of Y-family TLS polymerases to sites of DNA damage (via chaperoning, and by promoting PCNA mono-ubiquitylation), there is good evidence that Y-family polymerases can also associate with unmodified PCNA (i.e. non-ubiquitylated) via PIP boxes (72). In fact the extent to which PCNA monoubiquitylation is essential for TLS polymerase recruitment and lesion bypass has been heavily debated, with some investigators proposing that PCNA ubiquitylation is entirely dispensable for TLS (22,73–75). Third, the RAD18 and REV1/Polζ-mediated TLS sub-pathways appear to be separable and partially redundant (76). Thus, REV1-dependent TLS might compensate for RAD18-deficiency when PCNA is not mono-ubiquitylated.

Certain lesions, such as aflatoxin B1-induced DNA adducts, are known to result in G>T transversions, primarily due to the action of Polζ (77,78). Therefore, it is also possible that Rad18-independent TLS process mediated by Polζ contributes to SBS4 in 4NQO-treated mice. We also note that in our previous study, DMBA-induced A(T)>T(A) and G(C)>T(A) mutations also occurred to certain extent in the absence of Rad18. Therefore, TLS activities that are promoted by Rad18 may not necessarily depend absolutely on Rad18.

Interestingly, in this 4NQO-treatment study we observed the single bases substitution signature SBS39 exclusively in *Rad18*-deficient mice. Recent work suggests that SBS39 is associated with homologous recombination deficiency (79). Therefore, Rad18 may also shape cancer genomes via its established role in homologous recombination (80,81).

The divergent effects of Rad18 status on mutagenesis likely reflect the differential Rad18-dependencies of the TLS DNA polymerases that replicate damaged DNA, and whether those polymerases are error-free or error-prone for a given lesion. Mechanistically, the different Y-family DNA polymerases have different degrees of dependency on Rad18 for associating with PCNA at the replisome. For example, Polη has a relatively high-affinity PCNA-interacting peptide (PIP) box and can associate constitutively with the replisome, even when PCNA un-ubiquitylated (72,82,83). In contrast, the Polκ PIP box has low affinity for PCNA when compared with Polη, and may rely more heavily on Rad18-mediated PCNA ubiquitylation for its recruitment to the replisome (72). Thus, the error-propensity of RAD18-induced TLS will likely vary based on each specific combination of a Y-family DNA polymerase with its cognate or non-cognate DNA lesions.

In a previous study, Rad18 did not protect against UV-induced skin carcinogenesis (37), suggesting that error-free Polη-mediated bypass of UV-induced lesions (a tumor-suppressive activity) does not require Rad18. Mechanistically, Polη has a very high affinity PIP box which may be sufficient to mediate its PCNA binding even in the absence of Rad18-induced PCNA mono-ubiquitylation (72,82). Additionally, two other E3 ubiquitin ligases, HLTF and CRL4^Cdt2^, are capable of directly monoubiquitinating PCNA and promoting TLS in response to UV (70,71). Therefore, HLTF and CRL4^Cdt2^ might also compensate for Rad18-loss by sustaining Polη activity and suppressing UV-inducible skin carcinogenesis.

In a previous carcinogenesis experiment, we found that orally-administered DMBA induced several malignancies, including leukemia, spleen, liver and squamous skin carcinomas (36). Of those DMBA-induced malignancies, *Rad18* was tumor-suppressive for leukemia, and spleen and liver cancers, but not for skin carcinomas (which arose with similar incidence in WT and *Rad18^-/-^*mice). In that study, we only performed mutation signature analysis for the skin tumors (because these large clonal lesions were most amenable to genomic analysis).

For the DMBA-induced skin carcinomas, the genomic signatures were markedly different between *Rad18^+/+^* and *Rad18^-/-^*tumors. We found that *Rad18*-deficienct tumors had fewer A>T mutations, yet exhibited higher levels of >4bp deletions (36). We concluded that Rad18 promotes error-prone bypass of DMBA-adducted adenine residues. Moreover, in the absence of Rad18, DNA breaks caused by fork stalling most likely lead to deletion mutations (36). We did not identify the Rad18-dependent TLS polymerase(s) responsible for error-prone bypass of DMBA adducts. However, Polκ and Polη are good candidate mediators of Rad18 in DMBA-induced mutagenesis. Polκ performs replicative bypass of B[a]P-adducted dG and might also bypass the structurally-related DMBA-dG adducts. Furthermore, DMBA mutagenesis involves depurination of the adducted base followed by error-prone replication at the resulting AP site. Since Polη can perfom error-prone TLS at AP sites, this Y-family polymerase could also contribute to Rad18-dependent mutagenesis at DMBA-induced AP sites.

In contrast, the present study shows that Rad18 promotes error-free bypass of 4NQO-induced DNA lesions and suppresses 4NQO-induced oral carcinogenesis. Bulky C8- and N2- guanine and N6- adenine adducts are the primary DNA lesions induced by 4NQO. 4NQO is also known to generate oxidative stress-mediated 8-Oxo-7,8-dihydro-2’-deoxyguanosine (8-oxoG). Both types of DNA damage could lead to C>A transversions. The structural distortion of the guanine adduct can lead to polymerase-mediated erroneous mispairing with an adenine instead of cytosine. This leads to G:C>T:A transversions during a subsequent round of DNA replication. Similarly, 8-oxoG can also adopt a conformation that allows it to mispair with adenine and cause G:C>T:A transversions. RAD18 may prevent 4NQO-induced C>A transversions by recruiting a TLS polymerase that correctly introduces cytosine across from guanine bulky adducts. In this regard, Polκ is a good candidate effector of Rad18 since this polymerase is highly resilient to N2-guanine bulky adducts, is more efficient than Polη, and has preference equal to Polη for incorporating cytosine opposite N2-guanine adducts *in vitro* (84).

For error-free bypass of 4NQO-induced oxidative damage, Polη is a good candidate Rad18 effector which is highly efficient and accurate at incorporating cytosine against 8-oxoG (85).

Our transcriptomic analyses show that *Rad18* and *Polk* mRNAs were significantly induced in the oral epithelium from 4NQO-treated mice. This result is consistent with our hypothesis that Polη and Polκ mediate error-free bypass of lesions generated by 4NQO.

In addition to its role in mediating bypass of damaged DNA, RAD18-mediated TLS can also promote filling of ssDNA gaps such as those induced by oncogene-induced DNA replication stress RS in cancer cells (33,34). In our oncogene-induced lung tumorigenesis experiment, Rad18-deficiency did not significantly affect rates of *Kras^G12D^*-induced lung tumorigenesis or the mutational patterns of the *Kras^G12D^*-induced lung tumors (Supplementary Fig. 6). These findings may suggest that Rad18-dependent ssDNA gap filling (in cell harboring oncogene-induced RS) is relatively error-free. In many of our cell culture studies, genotoxin- or oncogene-induced DNA replication stress in RAD18 deficient cells led to aberrantly high levels of DSB markers such as pATM and γH2AX (33,36). Because DSBs can lead to structural variant (SV) formation, it is possible that 4NQO- or KRas-induced *Rad18^-/-^* tumors generated *in vivo* harbor increased numbers of SVs. However, the technical limitations of WES preclude scoring of the SV burden in the present study.

Because MAGE-A4 is a pathologically-relevant RAD18 activator in human cancers, it was also of interest to us to determine how MAGEA4 affects carcinogenesis in the 4NQO-induced tumorigenesis model. Many studies have suggested oncogenic roles for MAGEs and other CTAs, largely because these CTAs promote tumorigenic characteristics in tissue culture cells (86,87). Moreover, Armstrong et al. recently showed that MAGE-A4 expression mouse airway epithelia of *Pten*-deficient mice promotes NSCLC tumorigenesis (88).

Those workers proposed that MAGE-A4 stimulates endothelial cells to produce the cytokine CXCL12, which recruits immunosuppressive CXCR4+ plasma cells that inhibit T cell infiltration, thereby promoting tumor progression. In their study Armstrong et al. did not annotate mutation signatures and therefore it is not yet clear whether genome maintenance was affected by MAGE-A4 expression in the *Pten^-/-^* lung tumors.

In conclusion, we show that the important and extensively studied DNA repair factor Rad18 has tumor-suppressive activity. Given the prevalence of 4NQO-like chemical exposures to which humans are exposed, RAD18 is highly likely to shape human cancer genomes and perhaps influence other aspects of the tumorigenic process. However, the impact of RAD18 (and its pathological activator MAGE-A4) on mutagenic and tumorigenic outcomes is most probably highly context-specific and dependent on the specific chemically-induced DNA lesion, other genome maintenance factors (e.g. TLS polymerases), and perhaps other interacting genes.

## DATA AVAILABILITY

The raw RNA-seq data and the corresponding sample annotation file for control and 4NQO-treated mouse tongue epithelium are available under BioProject ID PRJNA1273114. The raw WES data for 4NQO-induced oral tumors from *Rad18^+/+^*, *Rad18^+/-^*, and *Rad18^-/-^* mice are available under BioProject ID PRJNA1273132. The raw WES data for 4NQO-induced oral tumors from MAGEA4-expressing (*Krt14-Cre*; *Rosa26^+/LSL-MAGEA4^)* mice and control *Rosa26^+/LSL-MAGEA4^* cohort are available under BioProject ID PRJNA1273474. The raw WES data for *Kras^G12D^*-induced lung tumors in *Rad18^+/+^* and *Rad18^-/-^* mice are available under BioProject ID PRJNA1273145. The Cancer Genome Atlas Head-Neck Squamous Cell Carcinoma (TCGA-HNSC) data was retrieved using TCGAbiolinks R package (version 2.30.0) and is available on TCGA portal (https://portal.gdc.cancer.gov/). Code used for analysis and visualization is available at https://github.com/JayRAnand/RAD18_mutagenesis and https://zenodo.org/records/15776968.

## SUPPLEMENTARY DATA

Supplementary Data is available at NAR online.

## ACKNOWLEDGEMENTS

We gratefully acknowledge the technical support from the UNC High Throughput Sequencing Facility. This facility is supported by the University Cancer Research Fund, Comprehensive Cancer Center Core Support grant (P30-CA016086) and the UNC Center for Mental Health and Susceptibility grant (P30-ES010126). We thank Nevaan Raval for assistance with The Cancer Genome Atlas (TCGA) data analysis and generating figures.

## Author contributions

Writing-Original Draft by J.R.A. and C.V.; Writing - Review & Editing by J.R.A., B.W., J.L., Q.G., Y.Y., G.D., D.W., A.J., Y.F., B.W., S.W., and C.V.; Investigation by J.R.A., B.W., Y.Y., G.D., D.W., A.J., Y.F., B.W.; Formal Analysis and Visualization by J.R.A., B.W., J.L., and Q.G.; Funding Acquisition by S.W. and C.V.; Methodology by J.R.A., B.W., J.L., Q.G., Y.Y., G.D., D.W., A.J., Y.F., B.W., S.W., and C.V.; Project Administration, Resources, Software, Data Curation by J.R.A., B.W., G.D., S.W., and C.V.; Supervision by S.W. and C.V..

