## Supplementary Figure for "Differential roles of Rad18 in repressing carcinogen- and oncogene-driven mutagenesis *in vivo*"

**Supplementary Figures**

**Figure S1: Heatmap of DDR gene mRNA expression in HNSCC patients from TCGA.**

**Figure S2. Gene ontology analysis of the 4NQO-induced transcriptome from preneoplastic oral epithelium**
(A) *Rad18*$^{+/+}$ mice were treated with 4NQO for 2 days or 14 days (or left untreated for controls). Then tongue epithelia from all three treatment groups were subjected to transcriptional profiling. A list of genes that were differentially expressed in the 4NQO treatment groups (with log fold change > |1| and a padj < 0.05 when compared with control mice) was inputted into the PANTHER analysis package. Statistically overrepresented biological processes identified by PANTHER are shown for the 2-day (left column) and 14-day (right column) 4NQO treatment groups. The orange arrowhead on the left column indicates activation of TLS genes in the 2-day 4NQO treatment group. (B) (D) Gene ontology (GO) analysis of common upregulated genes at day 2 and 14 in 4NQO-treated oral epithelium showing top 20 GO terms.

**Figure S3: Heatmap of DDR gene mRNA expression in 4NQO-treated mice (2 and 14 days).**

**Figure S4: Heatmap of DDR gene mRNA expression in 4NQO treated mice (study by *Lee et al.*).**

**Figure S5: RAD18 suppresses 4NQO-induced oral carcinogenesis.** (A) Survival curve showing that *Rad18*$^{-/-}$ (KO) mice develop tumors more rapidly than *Rad18*$^{+/+}$ (WT) littermates (*p* = 0.0338). Log-rank (Mantel-Cox) tests were performed to assess the statistical significance of survival curves. (B) Survival curve showing that *Rad18*$^{-/-}$ mice develop tumors more rapidly than *Rad18*$^{+/-}$ littermates (*p* = 0.0021). Log-rank (Mantel-Cox) tests were performed to assess the statistical significance of survival curves.

**Figure S6: RAD18 does not affect *Kras*-induced lung tumorigenesis.**
(A) Survival curve showing that *Rad18*$^{-/-}$ mice (n = 24) develop tumors at the same rate as Rad18-expressing littermates (n = 7) (p = 0.7356). Log-rank (Mantel-Cox) test was performed to assess the statistical significance of survival curves. (B) WES analysis of single nucleotide variants (SNV) in mouse lung tumors harvested from *Rad18*$^{-/-}$ (KO) and *Rad18*$^{+/+}$ (WT) animals. The data show that *Rad18*$^{-/-}$ tumors had similar levels of *Kras*$^{G12D}$-mediated mutations compared to *Rad18*$^{+/+}$ (p = 0.481). Statistical analysis to test the difference between two groups was performed using Wilcoxon rank sum test. (C) The data show that *Rad18*$^{-/-}$ tumors had similar transversions as *Rad18*$^{+/+}$ (padj < 0.001). Statistical analysis was performed using Wilcoxon test to test for the difference in number of SNVs between KO and WT samples at each level of transversions. (D) Statistical analysis of insertions and deletions (INDELs) in mouse oral tumors from the same samples and genotype groups as in (A). No statistically- significant differences were evident when comparing indel numbers between experimental groups except for 1bp-insertions (p = 0.00003). A two-way ANOVA was conducted to examine the effects of genotype (RAD18KO/WT) and mutational type (1bp insertions/deletions and 2-4bp insertions/deletions) on number of INDELs with a Bonferroni adjustment.

### Supplementary figures 1

## A

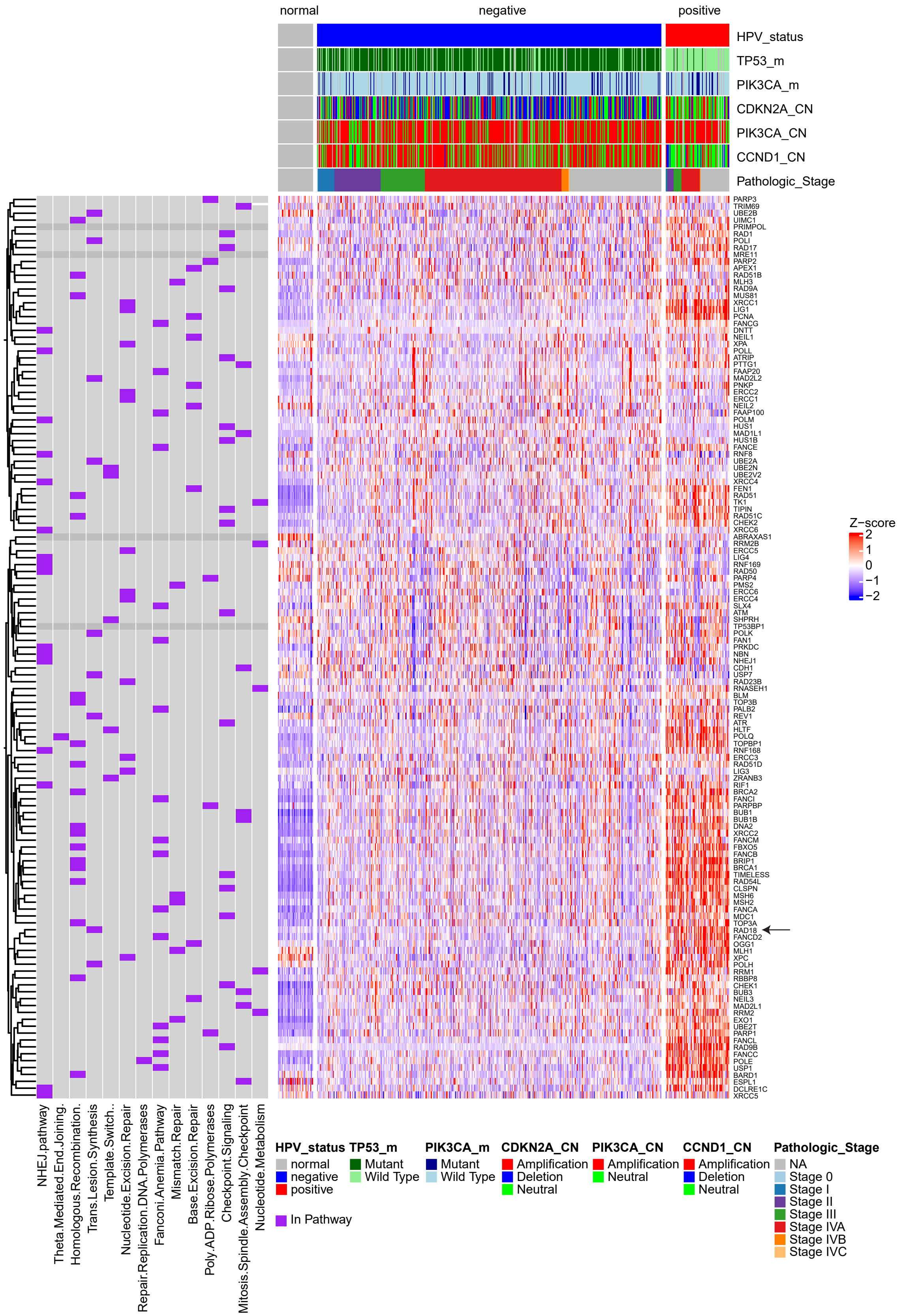

### Supplementary figures 2

A

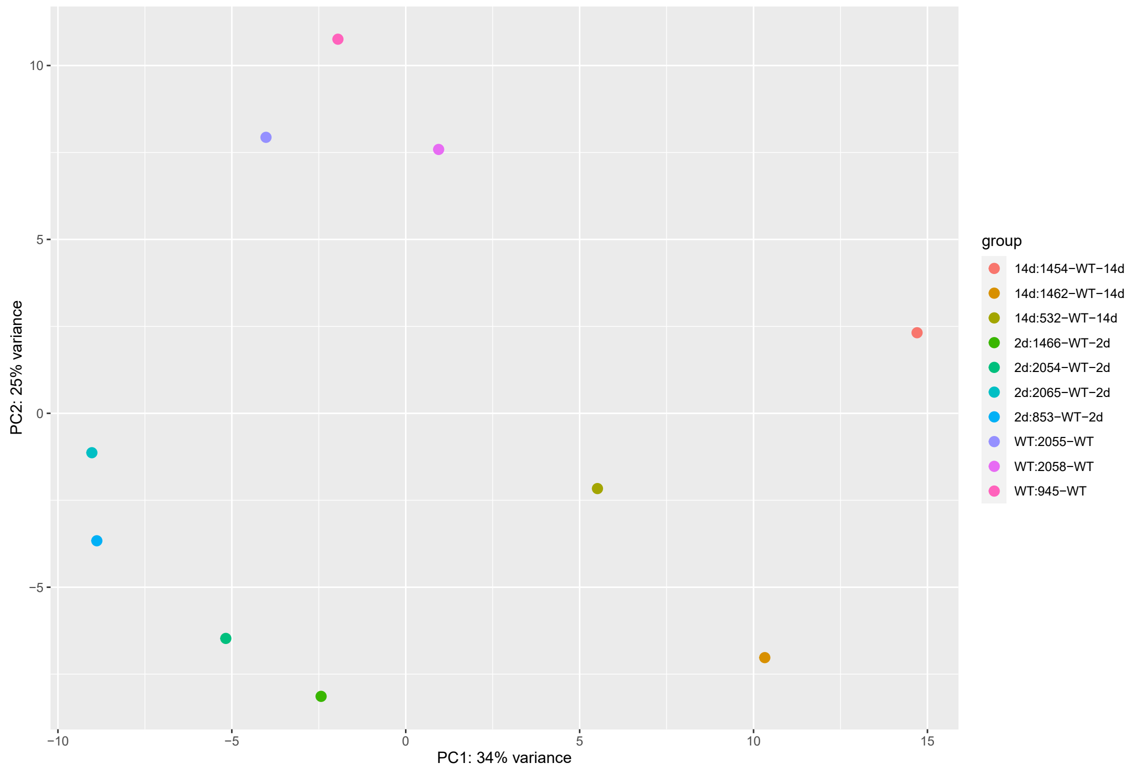


B

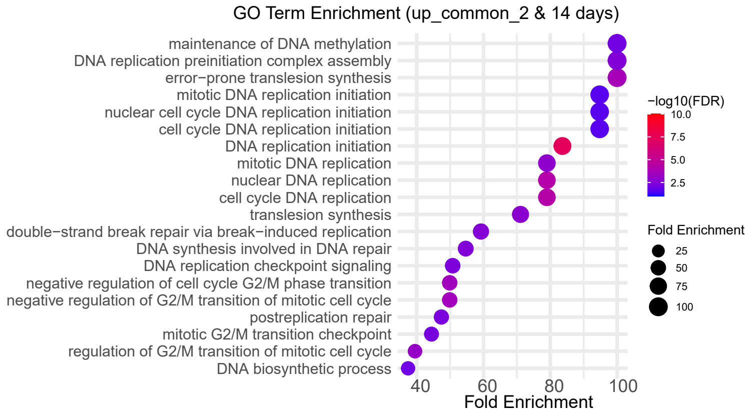

### Supplemnetary figures 3

## A

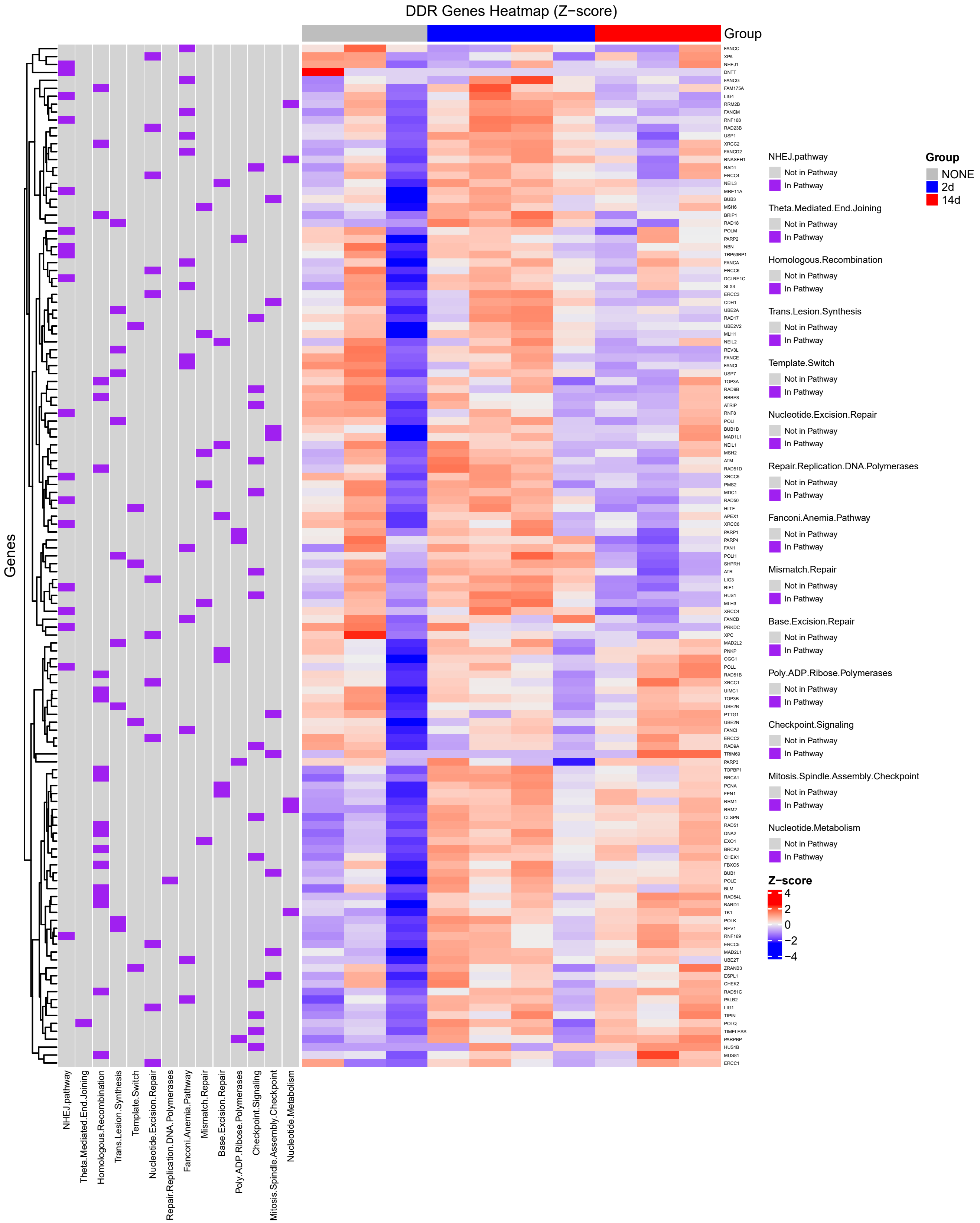

### Supplemnetary figures 4

## A

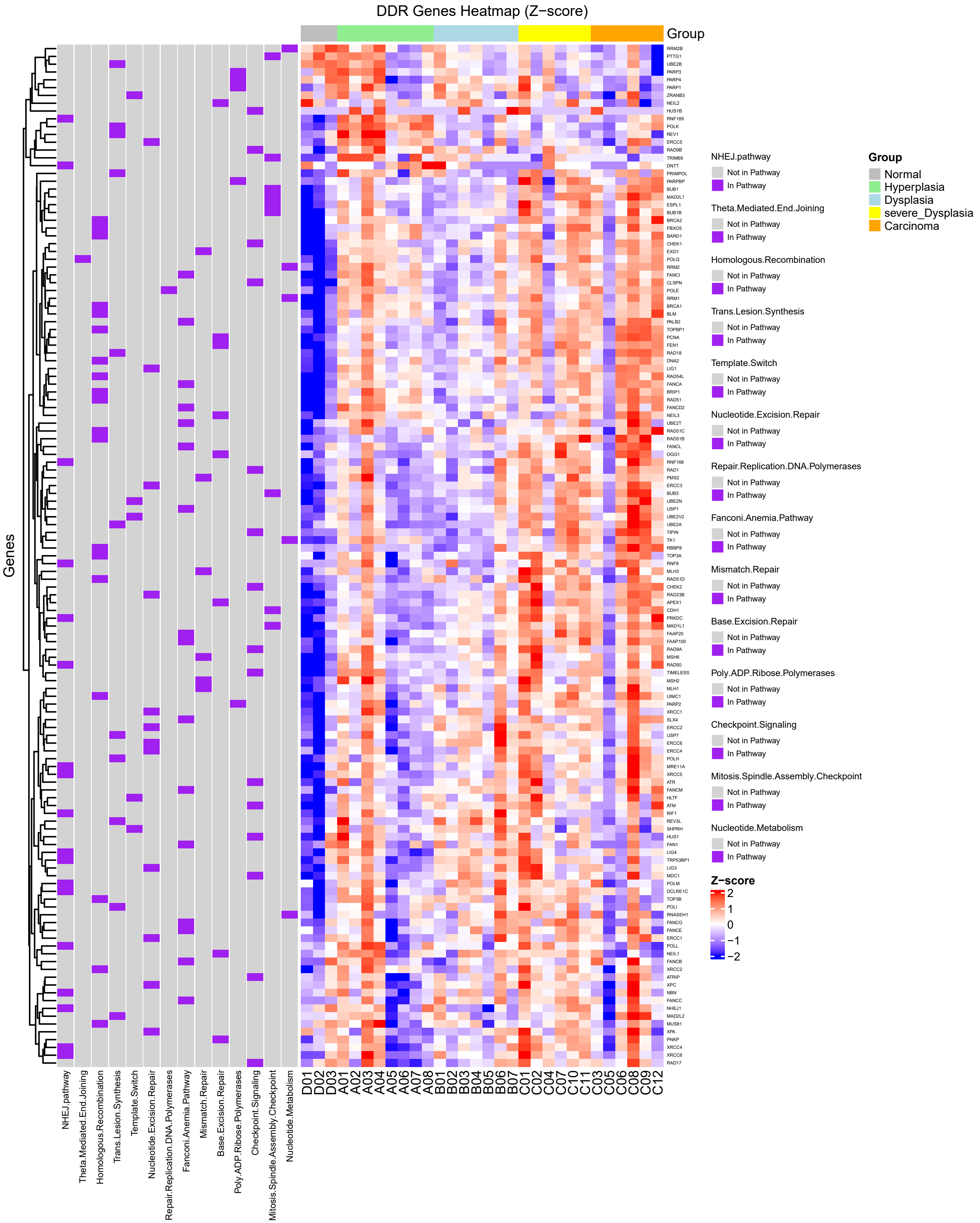

### Supplementary figures 5

A

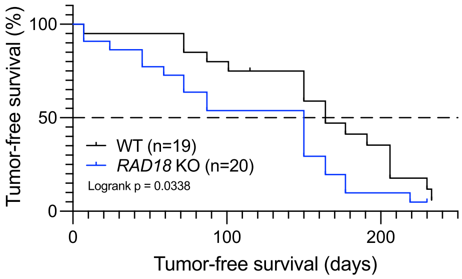


B

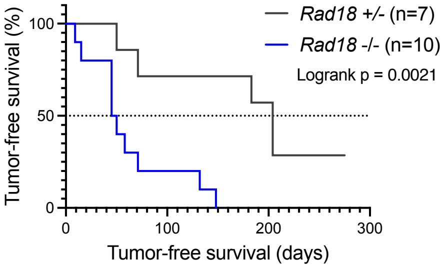

### Supplementary figures 6

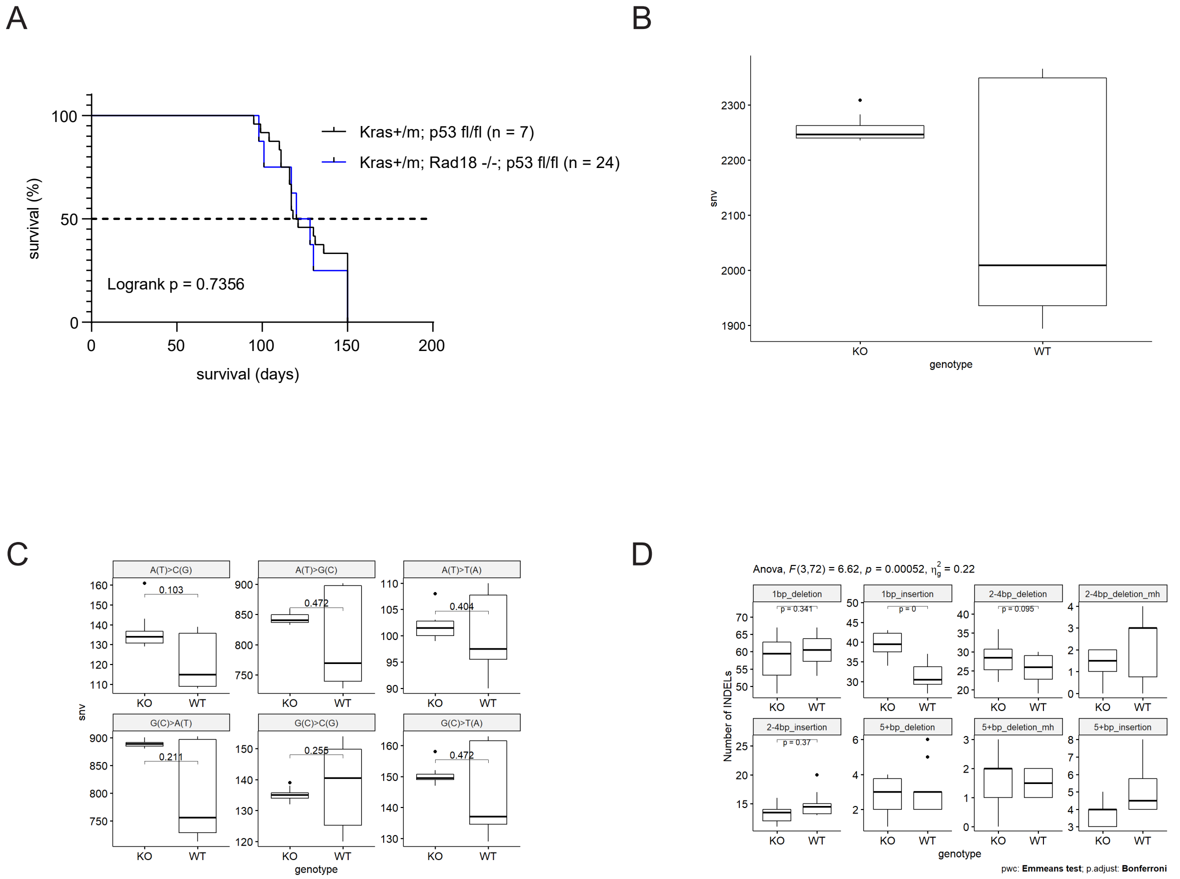
